# Distilling Trial-Level EEG Variability into Participant Representations: Neurophysiologically Constrained Graph Learning in ADHD

**DOI:** 10.64898/2026.10.01.755897

**Authors:** Rishabh Tiwari, Christian Beste, Negin Gholamipourbarogh

## Abstract

Extracting stable individual-level representations from noisy and variable trial-wise EEG remains a major challenge in computational cognitive neuroscience. We developed a spatiotemporal graph attention network to analyze Go/NoGo EEG from children and adolescents with and without ADHD. The framework combines ERP-informed temporal windows, phase-synchrony graph priors, and trial-to-participant transfer learning, making both information routing and representation geometry explicitly testable. The model learned reproducible attention trajectories, yet ADHD and control participants did not differ reliably in spatial attention allocation. Instead, group-related structure emerged in compact participant-level embeddings that distilled trial-level variability and retained discriminative information beyond convolutional representations. Performance exceeded a convolutional neural network comparison, but the central advance is methodological: neurophysiologically constrained graph learning can reveal where diagnostically relevant information emerges within the analytical pipeline. This provides a framework for developing auditable cognitive-neurophysiological markers of neurodevelopmental variation.

## 1. Introduction

Electroencephalography (EEG) is increasingly used in machine-learning approaches to patient stratification, symptom prediction, and diagnostic support. Yet, the central challenge is not only to maximize classification accuracy. EEG data are high-dimensional, noisy, and highly variable across individuals, recording sessions, and clinical subgroups. Under these conditions, flexible deep learning models may achieve apparently promising performance by exploiting regularities that correlate with diagnostic labels but are not mechanistically informative. Such shortcut learning [1] is particularly problematic in clinical neuroscience, where the goal is not only to predict group membership but also to identify representations that are robust, neurophysiologically plausible, and interpretable enough to support a mechanistic understanding of disorders [2–4].

Attention-Deficit/Hyperactivity Disorder (ADHD) provides a demanding test case for this problem [5]. ADHD is associated with impairments in cognitive control [6–8], including the ability to withhold inappropriate responses during response-inhibition tasks [9]. However, ADHD-related EEG signatures are unlikely to be captured by a single stationary marker or by a simple regional activation difference [10–12]. Inhibitory control unfolds over multiple processing stages [13], including early sensory selection, conflict monitoring, attentional updating, response cancellation, and later evaluative processes [14]. Moreover, ADHD has long been associated with increased intra-individual variability and altered state regulation [15–17], suggesting that clinically and mechanistically relevant information may reside not only in average event-related responses, but also in the organization of trial-to-trial variability. This creates a methodological problem. Event-related potential (ERP) approaches improve signal-to-noise ratio by averaging across trials [18,19], but this averaging necessarily suppresses the intra-individual variability that may be central to ADHD [16,20]. More recent multidimensional approaches have shown that ADHD-related response inhibition deficits can be reflected in temporally and spatially structured EEG activity profiles rather than in a single canonical component [20], fitting also recent advances in basic cognitive neuroscience research on response inhibition [21,22]. These findings indicate that diagnostically relevant neural information is distributed across multiple dimensions of the EEG signal, including time, space, and trial-to-trial variability. Consequently, patient-level classification models should be capable of learning these multidimensional patterns instead of relying on isolated temporal features or predefined electrophysiological markers.

To address this challenge, deep learning approaches have become increasingly attractive for EEG-based classification. Convolutional neural networks (CNNs) have emerged as a widely used architecture [23,24] because they can learn temporal and spatial filters directly from channel-by-time data [25]. However, CNNs primarily exploit local spatial neighborhoods defined by the convolutional kernel and do not explicitly model the functional interactions between distributed brain regions. Graph neural networks (GATs) offer an alternative by representing EEG sensors as nodes and their functional relationships as edges [26,27]. Within this framework, the focus extends beyond identifying informative temporal features at individual electrodes to learn how information is exchanged across the underlying brain network. This extension is important because cognitive operations depend on the transient integration and segregation of distributed neural populations [28,29]. Consequently, diagnostically relevant information may reside in the configuration of relationships among them. Oscillatory phase alignment provides a physiologically plausible basis for such coordination by regulating the temporal windows within which spatially separated neuronal populations can interact effectively [30]. Representing EEG recordings as connectivity-informed graphs therefore makes multivariate patterns of coordination available to the model. Such patterns are not captured when electrodes recording neural activity are considered primarily through their individual time series or spatial arrangement. This network-level perspective is particularly relevant to ADHD [31].

Therefore, we introduce a spatiotemporal GAT framework (ST-GAT) for patient-level ADHD classification from Go/NoGo EEG. Each trial is partitioned into ERP-informed temporal windows spanning early sensory processing, N2-related conflict monitoring, and P3-related motor inhibition [14]. For each ERP-informed temporal window, we used the phase-locking value (PLV) to define functional connectivity among electrodes, thereby creating a time-resolved graph structure: one PLV adjacency matrix per window, computed jointly over all trials and shared across the dataset. PLV was selected because it is well suited to capture oscillatory coordination among distributed neural populations [32,33], a property that is particularly relevant for response-inhibition tasks, where cognitive control emerges from dynamically evolving interactions between large-scale functional networks rather than isolated local components [14,34,35]. Beyond its neuroscientific relevance, PLV provides a natural graph representation of EEG activity by quantifying the strength of functional interactions between electrode pairs. The PLV yields bounded, symmetric, weighted adjacency matrices, making it directly compatible with graph-attention architectures in which edge strength constrains message passing between sensors. This raises the central methodological question of the present study: Does incorporating biologically motivated functional connectivity as a graph prior merely regularize a conventional deep-learning model, or does it enable the learning of qualitatively different patient-level representations? To address this question, we compare the proposed ST-GAT with the widely used EEGNet architecture [23] and examine performance, cross-seed stability, learned attention patterns, and the geometry of patient-level embeddings. We ask whether ADHD-related information is expressed as a group-specific spatial attention map, or whether it emerges in the way trial-level EEG variability is distilled into compact patient-level representations.

Our results support the latter account. The ST-GAT improves classification relative to EEGNet [23] across repeated cross-validation, but its most informative contribution lies in the learned representation space. The model recovers stable, time-resolved attention trajectories across ERP-relevant windows, yet ADHD and control trials do not show reliably different spatial attention allocations after correction for multiple comparisons. Instead, group separation is primarily expressed in the embedding geometry: while CNN-derived variability features contain widespread but redundant group differences, the ST-GAT compresses this information into a small set of non-redundant graph-derived dimensions. Additional decomposition analyses show that a substantial proportion of the discriminative GAT variance is not inherited from CNN features, indicating that the graph architecture captures spatiotemporal interaction structure that is missed by the baseline model. Together, these findings shift the focus of clinical EEG deep learning from regional biomarker localization to representation distillation. For ADHD, the relevant signal is not simply where the model attends, but how it transforms noisy trial-to-trial EEG dynamics into stable patient-level representations. The study shows how neurophysiologically constrained graph attention can make clinical EEG-AI more auditable: performance gains can be evaluated not only by accuracy, but by whether the model learns robust, sparse, and non-redundant representations that preserve meaningful structure in high-variability clinical data.

## 2. Methods

### 2.1. Participants and paradigm

Eighty-six children and adolescents (44 ADHD, 42 Control) participated in a visual Go/NoGo response-inhibition task. Participants pressed a button to Go stimuli (S11) and withheld their response to NoGo stimuli (S12). The ADHD group comprised clinically diagnosed patients recruited from the Department of Child and Adolescent Psychiatry, TU Dresden; controls were drawn from a community sample matched for age range. The groups differed significantly in age (ADHD: 11.1 ± 2.1 years; Control: 12.1 ± 2.3 years; t = −2.12, p = .037), IQ (ADHD: 99.7 ± 12.3; Control: 109.7 ± 10.5; t = −3.96, p = .00016), and sex distribution (ADHD: 84% male; Control: 36% male). Note that one of the ADHD participants had their information available for the EEG analysis but missing for this behavioral one.

### 2.2. EEG acquisition and preprocessing

EEG was recorded from 60 Ag/AgCl electrodes at an initial sampling rate of 500 Hz. Preprocessing was performed in two stages. The continuous data were first down-sampled to 256 Hz, band-pass filtered (0.5–40 Hz), average-referenced, and ocular-corrected via independent component analysis (ICA). The resulting preprocessed continuous recordings were exported for further processing. All recording files were verified to share identical preprocessing history (change sampling rate to 256 Hz; filters 0.5–40 Hz; new reference AVG; ocular correction; ICA). In Python (MNE-Python), the continuous data were epoched around the NoGo stimulus (S12) from −2.0 s to +2.5 s, baseline-corrected (−2.0 to 0 s), and subjected to a 200 µV peak-to-peak rejection criterion. NoGo trials were classified as Correct Rejections (CR; no button press within 1000 ms of the NoGo stimulus) or False Alarms (FA; button press within 1000 ms), Go trials were discarded. In the model pipeline, each preprocessed epoch was then cropped to 0.0–2.0 s post-stimulus (512 samples at 256 Hz), and channels were reordered to a standard 60-electrode montage. Per-channel z-scoring (mean and standard deviation computed on the training partition only) was applied within each cross-validation fold. No further filtering, re-referencing, or down-sampling was performed in Python. The final dataset comprised 8,237 NoGo epochs from 86 patients (44 ADHD: ∼4,010 epochs; 42 Control: ∼4,227 epochs), each with 60 channels and 512 time points.

### 2.3. Spatiotemporal graph construction

Each 2-s trial was partitioned into five ERP-informed temporal windows chosen to span the canonical processing stages of inhibitory control: W1, early sensory processing (0–200 ms, samples 0–51); W2, N2 conflict monitoring (200–350 ms, samples 51–90); W3, P3 motor inhibition (350–500 ms, samples 90–128); W4, late cognitive processing (500–1000 ms, samples 128–256); and W5, late slow cortical activity (1000–2000 ms, samples 256–512). Window boundaries were verified against the grand-average NoGo ERP: the N2 peaked at −6.2 µV at 324 ms over frontocentral electrodes (FCz, Fz, Cz, FC1, FC2), and the P3b peaked at +4.0 µV at 496 ms over centroparietal electrodes (Pz, CPz, CP1, CP2), confirming that each window captured its intended component. For each window, functional connectivity among the 60 electrodes was quantified using the phase-locking value (PLV) [32,33,36,37]. The analytic signal was obtained via the Hilbert transform, instantaneous phase was extracted, and the complex phase-coherence matrix was computed over all trials and time points within the window jointly (1):

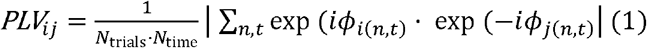

The resulting 60 × 60 PLV matrix (zero diagonal) was thresholded at the 80th percentile (top 20%), retaining 720 spatial edges per window. Across five windows, this yielded 720 × 5 = 3,600 spatiotemporal edges. Each edge carried a 6-dimensional attribute vector comprising the continuous PLV value and a one-hot encoding of window identity (5-d), enabling the graph-attention mechanism to condition message passing on both connection strength and temporal epoch. The spatiotemporal graph (edge index and edge attributes) was computed once on the full dataset before cross-validation. Because PLV is an unsupervised connectivity statistic computed without reference to diagnostic labels, this does not constitute label leakage. Nevertheless, the graph topology reflects the marginal phase-synchrony structure of the entire sample. This design choice was deliberate: the graph serves as a stable neurophysiological prior (constant across folds) rather than a fold-specific feature, ensuring that the model’s attention weights, not the graph structure, are the learned quantities evaluated across folds.

### 2.4. ST-GAT architecture

The ST-GAT comprises three components: a temporal convolutional backbone, a spatiotemporal graph-attention network, and a two-stage prediction head.

#### Temporal CNN backbone

A compact EEGNet-style convolutional network extracts a 256-dimensional per-trial embedding from the raw channel × time data. The backbone consists of: (i) a temporal convolution (1 → 8 channels, kernel 1 × 64, same padding), followed by BatchNorm and ELU activation; (ii) a depthwise spatial convolution (8 → 16 channels, kernel 60 × 1, 8 groups), followed by BatchNorm, ELU, and average pooling (factor 4); (iii) a depthwise-separable temporal convolution (16 → 16 channels, kernel 1 × 16, 16 groups), followed by a pointwise convolution (16 → 16, kernel 1 × 1), BatchNorm, ELU, average pooling (factor 8), and dropout (0.5). The output is flattened to a 256-dimensional trial embedding.

#### Node features

Each of the 60 graph nodes receives two contributions: (a) the 256-dimensional trial embedding broadcast across all nodes (identical at each electrode), and (b) a learned 60 × 256 node-identity embedding (one 256-dimensional vector per channel) that provides each electrode with a persistent positional prior. The per-node feature is the element-wise sum of these two contributions.

#### Spatiotemporal GAT

Two edge-conditioned graph attention layers (GATv2Conv; [38] process the 3,600-edge spatiotemporal graph. The first layer (256 → 64, 4 attention heads, concatenation → 256-dimensional) is followed by ELU. The second layer (256 → 64, 1 attention head) is followed by ELU. Edge attributes (6-dimensional) condition the attention coefficients, so that message passing is gated by both connection strength and temporal window.

#### Readout and pooling

A global attention readout (gate network: linear 64 → 32, ReLU, linear 32 → 1) aggregates the 60 node features into a single 64-dimensional graph-level vector per trial. The per-trial representation is the concatenation of the 256-dimensional CNN embedding and the 64-dimensional graph vector (320-dimensional).

#### Prediction heads

Two separate linear heads operate at different granularities. For trial-level pretraining, a linear head (320 → 1) produces a per-trial ADHD-vs-Control logit. For patient-level prediction, trials belonging to the same patient are aggregated via mean and standard deviation pooling across the 320-dimensional trial vectors, yielding a 640-dimensional patient embedding structured as [CNN_mean_ (256), GAT_mean_ (64), CNN_std_ (256), GAT_std_ (64)]. A patient head (linear 640 → 64, ReLU, dropout 0.3, linear 64 → 1) produces the patient-level logit.

Figure 1 illustrates the full pipeline (see methods section): EEG epoch → temporal window partitioning → per-window PLV graph (720 edges each) → ST-GAT processing → trial-level embeddings → patient-level aggregation → ADHD/Control classification.

**Figure 1.**
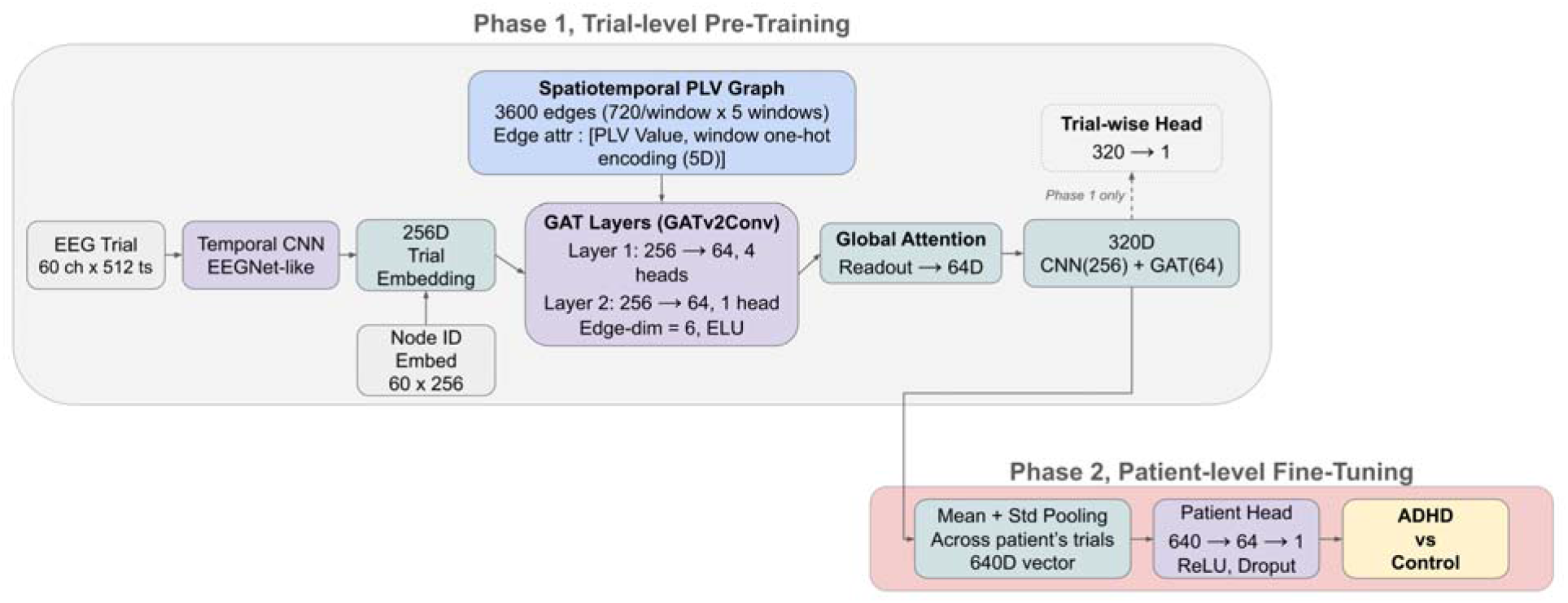
Architecture and two-phase training workflow of the spatiotemporal graph attention network (ST-GAT). During Phase 1, an EEGNet-style CNN generates a 256-dimensional trial embedding, which is combined with learned node identities across 60 sensors. Five ERP-informed PLV graphs constrain spatiotemporal message passing through two GATv2 layers, producing a 64-dimensional graph representation. This representation is concatenated with the CNN embedding into a 320-dimensional trial vector for classification and pretraining. During Phase 2, mean and standard-deviation pooling across trials yields a 640-dimensional patient-level embedding for ADHD vs control classification. During fine-tuning, the CNN backbone remains frozen while the graph-attention components and patient-level classifier are optimized. *Abbreviations:* ch, channels; ts, time samples; d, dimensions; h, attention heads; PLV, phase-locking value; CNN, convolutional neural network; GAT, graph attention network.

### 2.5. Transfer learning and training

Training follows a two-phase transfer-learning protocol. Both phases are nested within each outer cross-validation fold, ensuring that the backbone is never reused across outer folds.

#### Phase 1: trial-level pretraining

The backbone (CNN, node-identity embedding, GAT, readout) and trial head are jointly optimized for ADHD-vs-Control classification at the trial level on NoGo-ERP epochs. A 5-fold inner stratified group k-fold (patient-group-aware) selects the best backbone by minimum validation loss. The loss is binary cross-entropy with logits, with class-imbalanced positive weight computed from the training labels. Optimization uses Adam (learning rate 1 × 10⁻³, weight decay 1 × 10⁻) for 15 epochs, batch size 64. An attention-entropy term (λ = 0.01) is logged as a sparsity diagnostic; it is computed without gradient tracking and does not contribute to gradient updates.

#### Phase 2: patient-level fine-tuning

The best Phase 1 backbone is loaded and frozen: all convolutional, batch-normalization, and depth-wise parameters have their gradients disabled, and BatchNorm layers are held in evaluation mode. The GAT layers, readout, node-identity embedding, and patient head remain trainable. A 5-fold inner stratified k-fold (patient-level) isolates a validation set. Optimization uses AdamW with differential learning rates: the patient head is trained at 5 × 10⁻ (weight decay 1 × 10⁻²), while the GAT layers, readout, and node-identity embedding are trained at 1 × 10⁻ (weight decay 1 × 10⁻). Training proceeds for up to 15 epochs with early stopping (patience = 5, monitored on validation accuracy), batch size 1 (one patient per batch). The loss is binary cross-entropy with logits, with positive weight recomputed from the Phase 2 training labels. At inference, the five surviving Phase 2 models (one per inner fold) are ensembled: their sigmoid probabilities are averaged, and a threshold of 0.5 yields the binary prediction for each test patient. The patient-level embedding (640-dimensional) is extracted from the final Phase 2 model for downstream interpretability and embedding analysis.

### 2.6. EEGNet baseline

To isolate the contribution of the graph-attention component, an EEGNet-style baseline was trained under an identical protocol. The baseline shares the same temporal CNN backbone as the ST-GAT (same kernel sizes, filter counts, normalization, activation, pooling, and dropout), but includes no graph structure, no GAT layers, and no node-identity embedding. The per-trial embedding (256-dimensional) is aggregated to the patient level via mean and standard deviation pooling, yielding a 512-dimensional patient vector. The same two-phase transfer-learning scheme, cross-validation scheme, and hyperparameters are applied, with the obvious adjustment that Phase 2 fine-tuning optimizes only the patient head (learning rate 5 × 10⁻, weight decay 1 × 10⁻²), as there are no GAT parameters to update.

### 2.7. Evaluation and statistics

Model performance was estimated via 5 seeds (42, 101, 123, 456, 789) × 5-fold patient-level stratified k-fold cross-validation (outer loop, n_test_ ≈ 17–18 per fold), yielding 25 test-set evaluations per model. Per-seed accuracy was computed as the ensemble accuracy across the five outer folds. Sensitivity, specificity, balanced accuracy, and area under the ROC curve (AUC) were computed per seed from the saved per-patient probabilities. The ST-GAT and EEGNet baseline were compared via a paired t-test across the five seed-level accuracies. The seed yielding the lowest accuracy for both models (seed 789) was reported as a caveat; performance excluding this seed was also reported.

#### Embedding analyses

Patient-level embeddings (640-dimensional for ST-GAT, 512-dimensional for EEGNet) were extracted from the final Phase 2 model of each outer fold. Four component blocks were analysed separately: CNN_mean_ (256-d), GAT_mean_ (64-d), CNN_std_ (256-d), and GAT_std_ (64-d). Univariate group differences were tested per dimension via Mann-Whitney U with Bonferroni correction (α / dimensions). Multivariate group separation was tested via Hotelling’s T²; for blocks exceeding 100 dimensions, principal component analysis (PCA) was applied to retain 90% of variance before the T² test, and the effective rank (k_PCA_) was reported.

#### Variance decomposition (MANCOVA)

To determine whether the GAT-derived embeddings carry discriminative information beyond what a CNN captures, a multivariate analysis of covariance (MANCOVA) was performed using Wilks’ Λ. Two covariate framings were reported: (a) the independent EEGNet baseline embeddings (the stronger test, asking whether the GAT contributes beyond a separately trained CNN), and (b) the internal CNN branch of the ST-GAT itself (testing whether the GAT adds signal beyond its own backbone). In both cases, covariates were PCA-reduced to 90% variance. Effect-size decomposition was quantified as the proportion of the GAT’s multivariate Cohen’s d² remaining after residualisation (GAT-unique %) versus absorbed by the covariates (CNN-inherited %). This decomposition is invariant to the test statistic. Reporting of standardized effect sizes alongside null-hypothesis tests adheres to COBIDAS MEEG guidelines [39].

### 2.8. Interpretability analyses

Spatial attention was aggregated at four levels of granularity: 60 individual channels, 12 sub-regions, 5 lobes, and 25 lobe-pair connectivity profiles, using the canonical sub-region → lobe mapping (centroparietal → parietal, frontotemporal → frontal, temporoparietal → parietal). Per-window attention trajectories were computed as the lobe-level proportion of total attention within each ERP window, averaged across the five seeds. Patient-level embeddings were visualized via t-SNE, with PCA pre-reduction to 30 dimensions for computational efficiency, following established guidelines for high-dimensional biological data [40]. Cross-method interpretability via GNNExplainer, SHAP, and Grad-CAM was not performed and is listed as a future direction.

## 3. Results

### 3.1. Spatiotemporal graph construction creates a neurophysiologically constrained representation space

The ST-GAT operates on a spatiotemporal graph that encodes phase-synchrony structure across five ERP-informed windows. The graph imposes a neurophysiological prior that constrains message passing to empirically observed phase-synchrony connections, rather than allowing unconstrained spatial filtering. This design is not merely a classification architecture but a structured representation-learning framework in which the graph prior biases the model toward oscillatory coordination patterns relevant to response inhibition. Figure 2 shows five temporal windows spanning the canonical processing stages (early sensory, N2, P3, late, slow cortical), and window boundaries were verified against the grand-average NoGo ERP (N2 at −6.2 µV / 324 ms; P3 at +4.0 µV / 496 ms).

**Figure 2.**
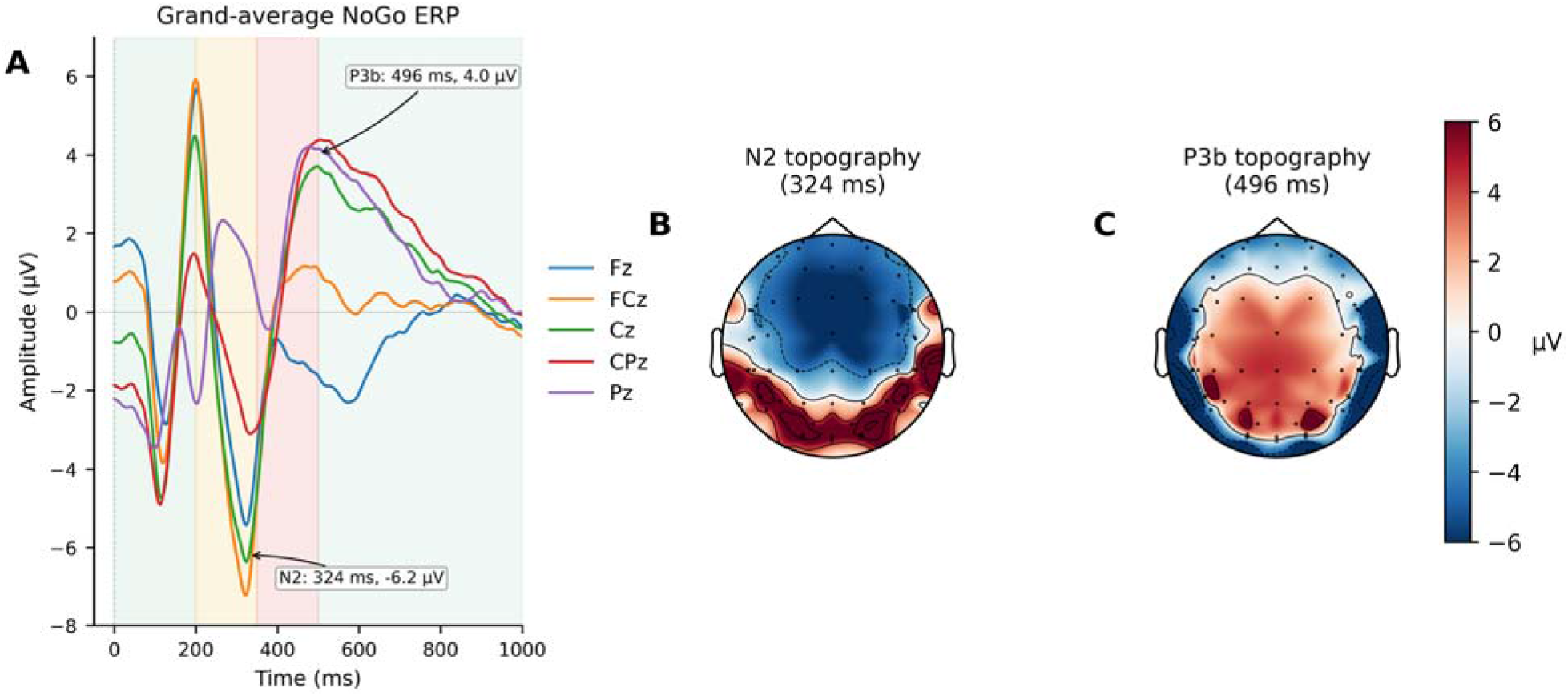
Grand-average NoGo event-related potential (ERP) waveforms and scalp topographies validating the ERP-informed temporal segmentation. **(A)** Grand-average NoGo ERP waveforms at selected frontocentral and centroparietal electrodes (Fz, FCz, Cz, CPz, Pz). Shaded background bands indicate the five ERP-informed temporal windows: Early (0–200 ms), N2 (200–350 ms), P3 (350–500 ms), Late (500–1000 ms), and Slow Wave (1000–2000 ms). Arrows mark the frontocentral N2 peak (−6.2 µV at 324 ms) and the centroparietal P3b peak (+4.0 µV at 496 ms). **(B and C)** Scalp topographies at the N2 and P3b peak latencies confirm the expected frontocentral N2 and centroparietal P3b distributions, supporting the window placement used for the spatiotemporal graph construction. *Abbreviations:* ERP, event-related potential; NoGo, no-go.

### 3.2. ST-GAT improves patient-level ADHD classification relative to EEGNet

Table 1 reports per-seed performance for both models across the 5-seed × 5-fold patient-level cross-validation. The ST-GAT achieved a mean accuracy of 75.3% ± 6.34% (range 62.8– 80.2%), compared to 70.0% ± 3.92% (range 62.8–74.4%) for the EEGNet baseline, corresponding to a +5.3 percentage-point advantage. Sensitivity, specificity, balanced accuracy, and AUC were consistently higher for the ST-GAT across most seeds (Table 1; AUC range: ST-GAT 0.68–0.84, EEGNet 0.74–0.83).

**Table 1.** Patient-level classification performance of ST-GAT and EEGNet across cross-validation seeds.

| Seed | ST-GAT |  |  |  |  | EEGNet |  |  |  |  |
| --- | --- | --- | --- | --- | --- | --- | --- | --- | --- | --- |
|  | Accuracy | Sensitivity | Specificity | Bal-Acc | AUC | Accuracy | Sensitivity | Specificity | Bal-Acc | AUC |
| 42 | **77.90%** | 75.00% | **81.00%** | **78.00%** | **0.843** | 72.10% | 75.00% | 69.00% | 72.00% | 0.813 |
| 101 | **77.90%** | **75.00%** | **81.00%** | **78.00%** | **0.825** | 69.80% | 65.90% | 73.80% | 69.90% | 0.782 |
| 123 | **80.20%** | **75.00%** | **85.70%** | **80.40%** | **0.84** | 74.40% | 72.70% | 76.20% | 74.50% | 0.818 |
| 456 | **77.90%** | **75.00%** | **81.00%** | **78.00%** | 0.783 | 70.90% | 68.20% | 73.80% | 71.00% | **0.825** |
| 789 | 62.80% | **59.10%** | 66.70% | 62.90% | 0.681 | 62.80% | 56.80% | **69.00%** | 62.90% | **0.745** |

**Note:** Balanced accuracy (Bal-Acc) is the arithmetic mean of sensitivity and specificity. All values except AUC are expressed as percentages. Bold values indicate the better-performing model for the respective seed and metric; ties are not highlighted. **Abbreviations:** ST-GAT, spatiotemporal graph attention network; Bal-Acc, balanced accuracy; AUC, area under the receiver-operating-characteristic curve.

Seed 789 yielded the lowest accuracy for both models (∼62.8%), suggesting an unfavorable patient split rather than a model-specific failure. Excluding this seed, the ST-GAT reached 78.5% ± 1.0% versus 71.8% ± 1.7% for EEGNet (Δ = +6.7 pp), demonstrating that the performance gain is not driven by a single lucky split. The paired seed-wise plot (Figure 3) visually confirms that the ST-GAT advantage holds across the remaining four seeds, with the two models converging only at the outlier seed.

**Figure 3.**
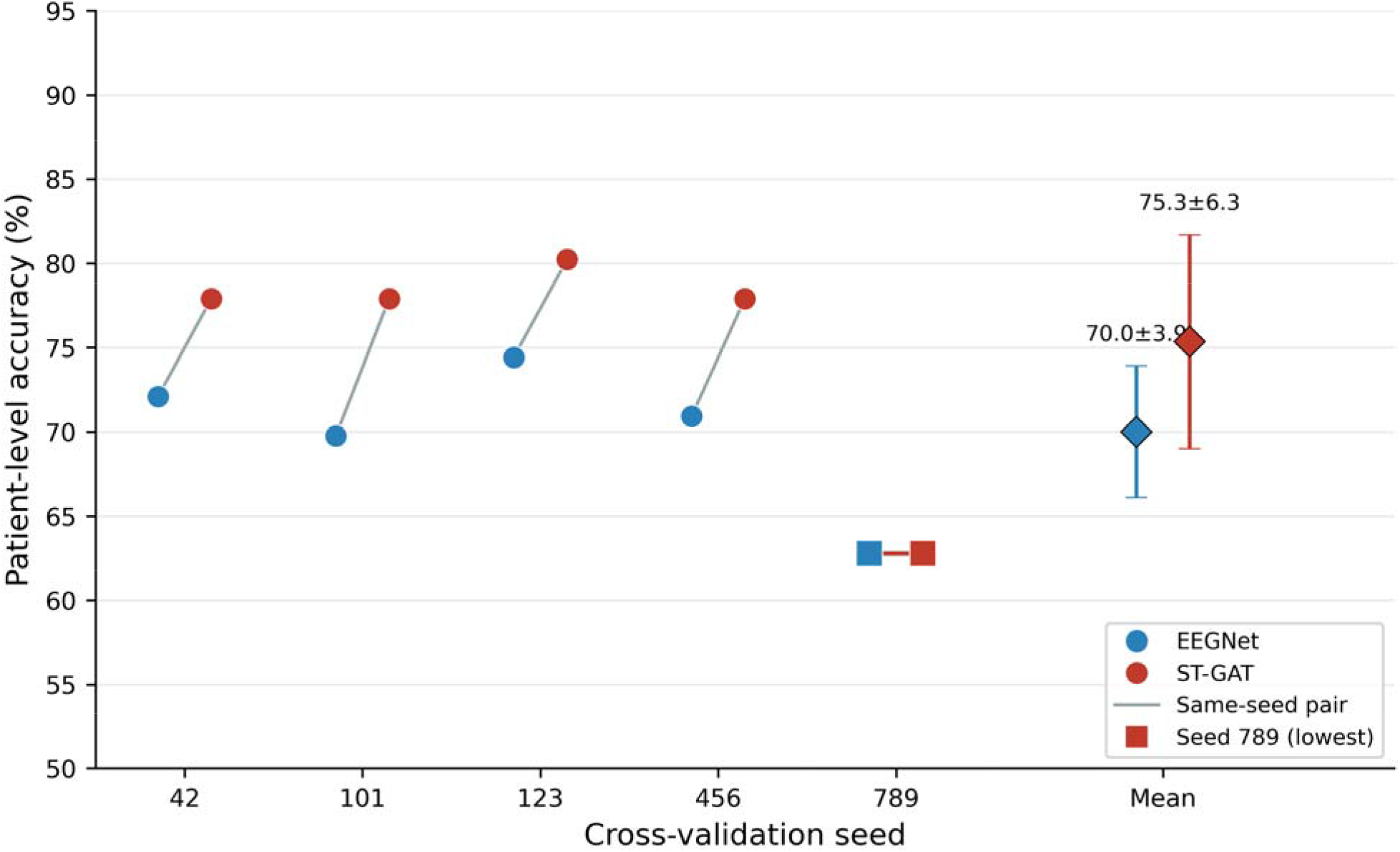
Patient-level classification accuracy of ST-GAT and EEGNet across repeated cross-validation seeds. Points represent patient-level accuracies obtained for each of five random seeds after combining the held-out predictions from the five outer patient-level cross-validation folds. Grey lines connect results obtained from the same seed, thereby showing the paired comparison between EEGNet and ST-GAT. Blue and red circles denote EEGNet and ST-GAT accuracies, respectively. Points with CI to the right indicate the mean ± SD across all five seeds. Red squares and the red connecting line identify seed 789, which produced the lowest accuracy for both architectures and was retained in the primary analysis. *Abbreviations*: ST-GAT, spatiotemporal graph attention network; SD, standard deviation; CI, confidence interval.

### 3.3. Learned attention patterns are stable across seeds

To establish that the model learns reproducible spatiotemporal structure rather than seed-dependent patterns, cross-seed attention stability was assessed at four spatial granularities via pairwise Spearman ρ within each window. Channel-level stability was high (ρ = 0.88 ± 0.06), and sub-region stability was even higher (ρ = 0.90 ± 0.12), indicating that broad spatial allocation patterns are highly reproducible. Lobe-level stability was moderate (ρ = 0.78 ± 0.21), reflecting greater variability at coarser aggregation. Lobe-connectivity stability was exceptional (ρ = 0.98 ± 0.01), indicating that the model’s routing profile over the 25 lobe-pair connections is nearly identical across seeds. Friedman tests confirmed non-uniform attention allocation at the lobe-connectivity level (χ² = 36.56, d_f_ = 24, p = 2.22 × 10), while lobe-level allocation did not deviate significantly from uniformity across seeds (χ² = 0.32, d_f_ = 4, p = .988).

### 3.4. ST-GAT recovers a task-relevant temporal attention trajectory

The model’s window-resolved attention allocation was examined across the five ERP windows (Figure 4). Figure 5 reveals that attention was broadly shared between frontal and parietal regions across all windows, with parietal attention peaking in the P3 time window (W3: 44.4% parietal, 35.5% frontal). This pattern is consistent with the known involvement of centroparietal regions in P3 generation. The top channels in W3 were Pz (2.92% of window attention) and CP2 (2.38%), both of which appeared in the top-3 channels across all 5 seeds (Pz: 5/5 seeds; CP2: 5/5 seeds), confirming stable parietal P3b recovery.

**Figure 4.**
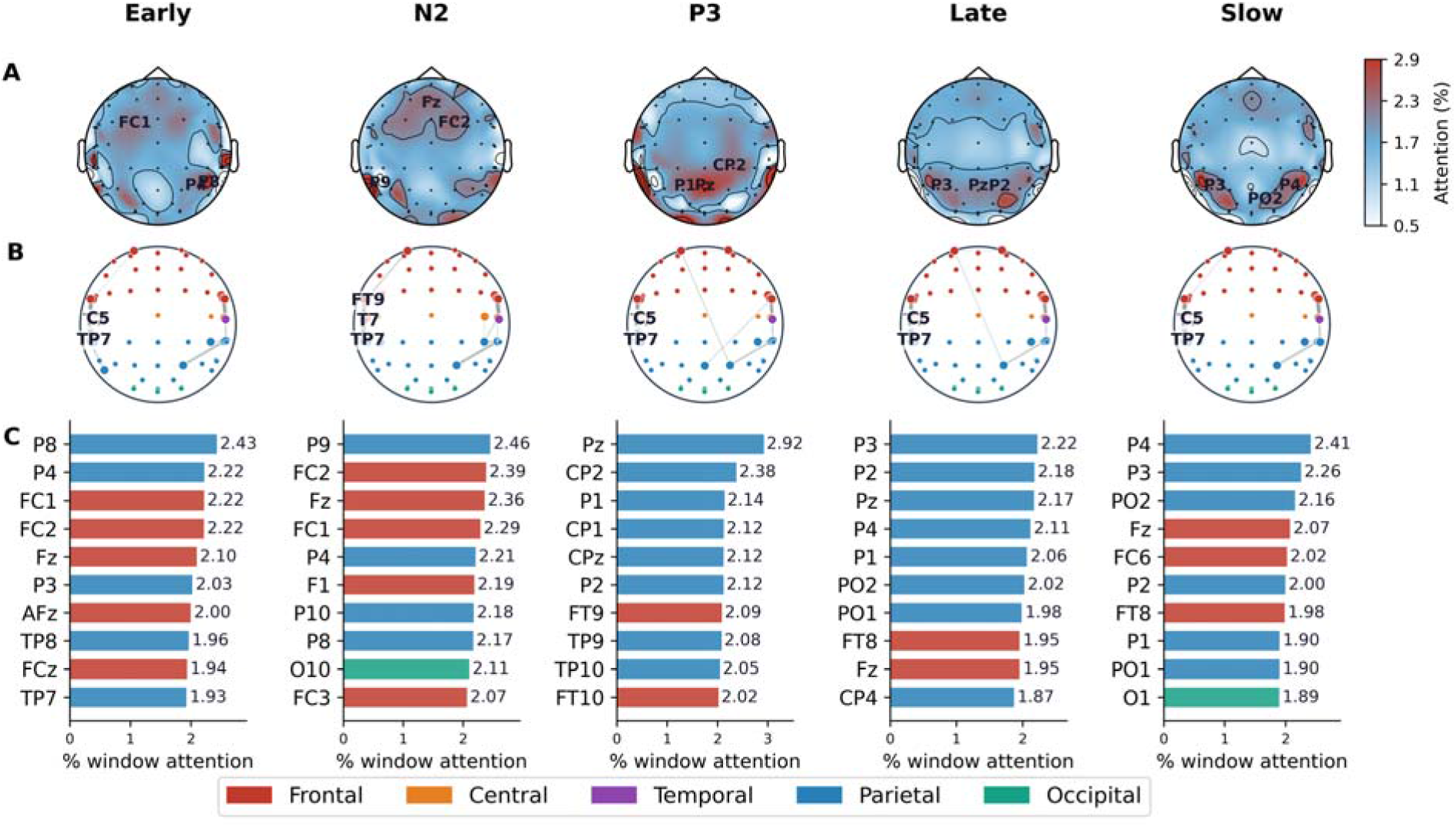
Window-resolved spatial and connectivity attention profiles learned by the ST-GAT. **(A)** Scalp topographies show the fraction of total channel attention allocated within each temporal window. Electrode positions are indicated by circles, and the five channels receiving the closest attention in each window are labeled. **(B)** Circular connectivity plots show the 30 highest-ranked graph-attention edges within each window on the PLV-constrained sensor graph. Electrode positions are arranged schematically and colored according to their assigned lobe. **(C)** Bar plots show the ten channels receiving the highest proportion of total attention in each window; numerical labels indicate the percentage of window-specific attention. Colors identify frontal (red), central (orange), temporal (purple), parietal (blue), and occipital (green) electrodes. The displayed attention values represent model-derived routing weights on the PLV-defined graph and should not be interpreted as raw PLV connectivity strength. *Abbreviations:* ERP, event-related potential; PLV, phase-locking value; ST-GAT, spatiotemporal graph attention network.

**Figure 5.**
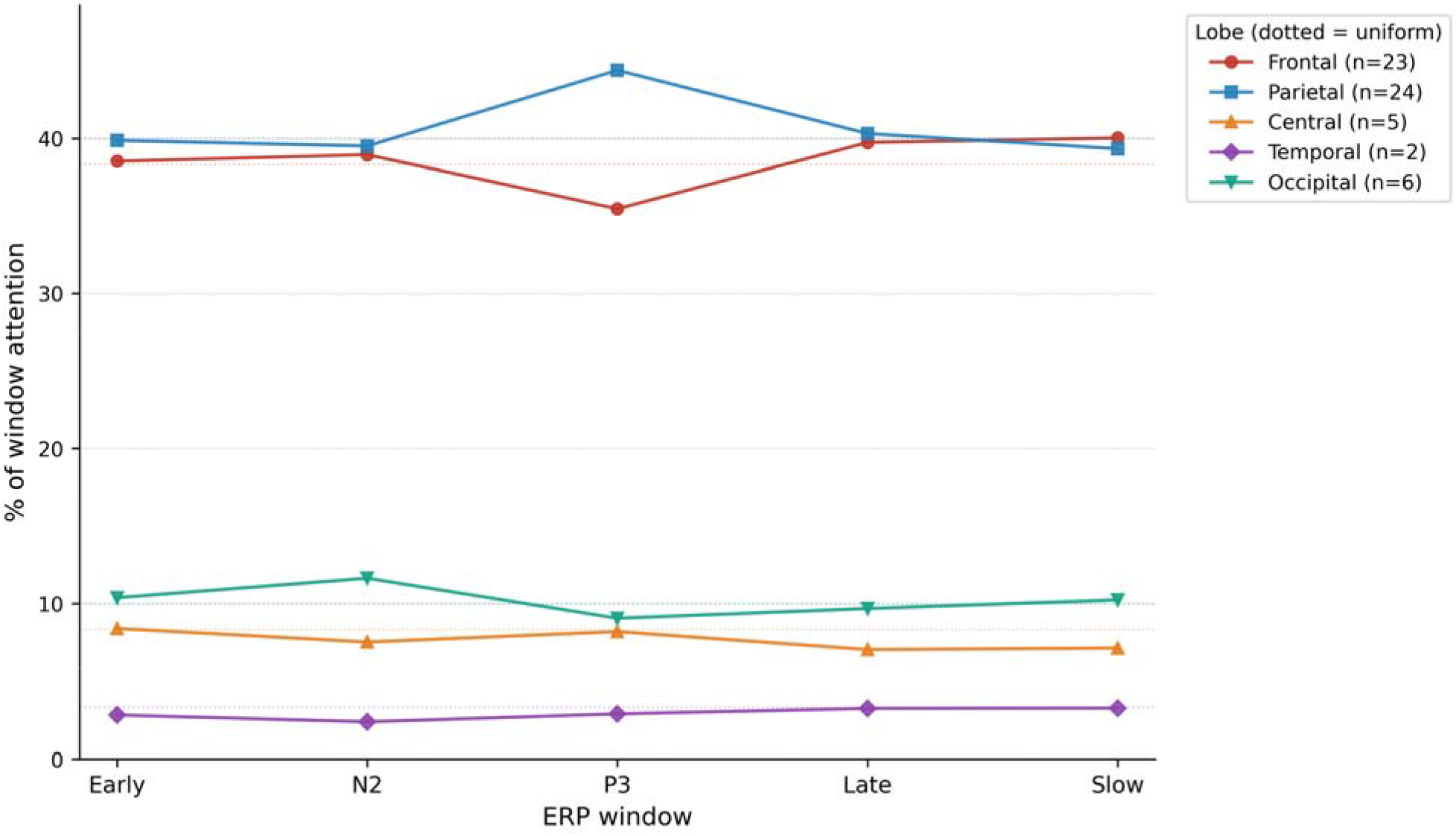
Lobe-level attention trajectories across the five ERP-informed temporal windows. Each line shows the percentage of total window attention allocated to electrodes within a given lobe, averaged across the five random seeds (n = 44 ADHD, n = 42 controls). Dotted horizontal lines indicate the uniform-attention reference corresponding to the proportion of electrodes in each lobe. The trajectories reveal reproducible routing preferences across windows: frontal and central attention dominate early and N2 windows, parietal attention rises during the P3 and late windows, and occipital attention remains relatively low throughout. *Abbreviations:* ERP, event-related potential.

In the early window (W1), frontal and parietal attention were co-dominant (38.5% and 39.9%, respectively), with P8 (5/5 seeds) and P4 (3/5 seeds) as the most consistent top channels. The N200 window (W2) showed a similar frontal–parietal profile (38.9% / 39.5%), with P9 (5/5 seeds) and FC2 (4/5 seeds) as the most consistent channels. The late windows (W4, W5) maintained frontal–parietal co-dominance without strong shifts to other regions. The overall trajectory thus reflects a stable, task-relevant spatiotemporal routing profile spanning frontocentral and centroparietal regions across the inhibitory-control processing cascade.

### 3.5. ADHD and control trials do not differ in spatial attention allocation

A central question was whether the model classifies ADHD by attending to different scalp regions in patients versus controls. Per-class attention weights were compared across 25 lobe × window combinations via Mann-Whitney U with Bonferroni correction (α = 0.002). No comparison survived correction (0/25). The maximum absolute effect size was |r| = 0.15. ADHD (220 patient-fold-instances) and Control (210 patient-fold-instances) attention allocations were thus statistically indistinguishable at the spatial level after correction.

This null result is not a weakness but a pivotal finding: it indicates that the model does not solve the classification problem via a group-specific spatial attention map. Both groups appear to engage a broadly similar task-relevant sensor-level architecture, while diagnostic information emerges in the geometry of the patient-level representations derived from this architecture.

### 3.6. Patient-level embeddings reveal that ADHD information lies in trial-to-trial variability

To identify where diagnostic information resides in the representation space, the four component blocks of the 640-dimensional patient embedding were analysed separately: CNN_mean_ (256-d), GAT_mean_ (64-d), CNN_std_ (256-d), and GAT_std_ (64-d).

Univariate analysis (Mann-Whitney U per dimension, Bonferroni-corrected) revealed a marked asymmetry between mean and variability features (Table 2). The CNN_std_ block contained the highest proportion of individually significant dimensions (248/256 = 97%), consistent with the hypothesis that trial-to-trial variability carries widespread group-discriminative signal. However, the discriminative structure of CNN_std_ was low-rank: PCA retained only 28 components to explain 90% of variance, indicating that the 248 significant dimensions are highly correlated and largely redundant. In contrast, GAT_std_ showed far fewer individually significant dimensions (12/64 = 19%) yet achieved the strongest multivariate separation, indicating that the graph-attention mechanism compresses diffuse variability into a compact, non-redundant representation. The GAT_mean_ block (9/64 = 14% Bonferroni-significant) and CNN_mean_ block (43/256 = 17%) carried comparatively sparse univariate signal.

**Table 2.**
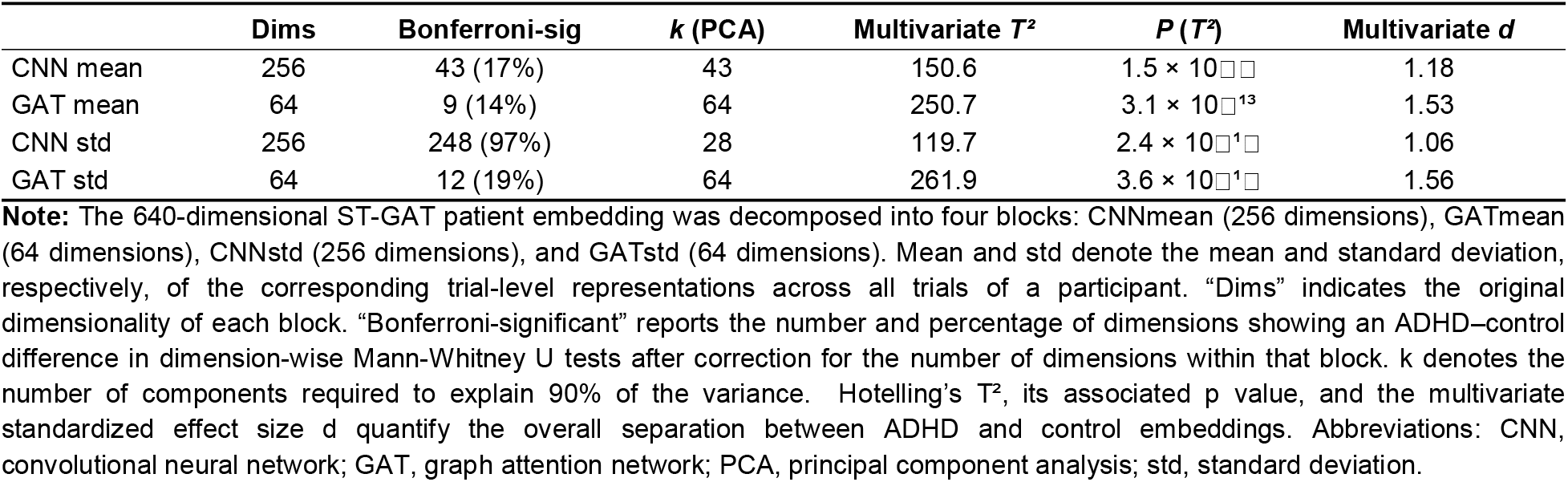
Univariate and multivariate ADHD–control separation within the patient-level embedding components.

Multivariate analysis (Hotelling’s T², with PCA pre-reduction for blocks > 100 dimensions) confirmed that all four blocks exhibit significant ADHD-vs-Control separation. The GAT blocks achieved the strongest multivariate effect sizes: GAT_std_ (T² = 261.9, d = 1.56, p = 3.6 × 10 ¹) and GAT_mean_ (T² = 250.7, d = 1.53, p = 3.1 × 10 ¹³) substantially exceeded CNN_std_ (T² = 119.7, d = 1.06) and CNN_mean_ (T² = 150.6, d = 1.18). Patient-level t-SNE visualisation (Figure 6) illustrates the multivariate group separation in the full 640-d space and within each sub-block.

**Figure 6.**
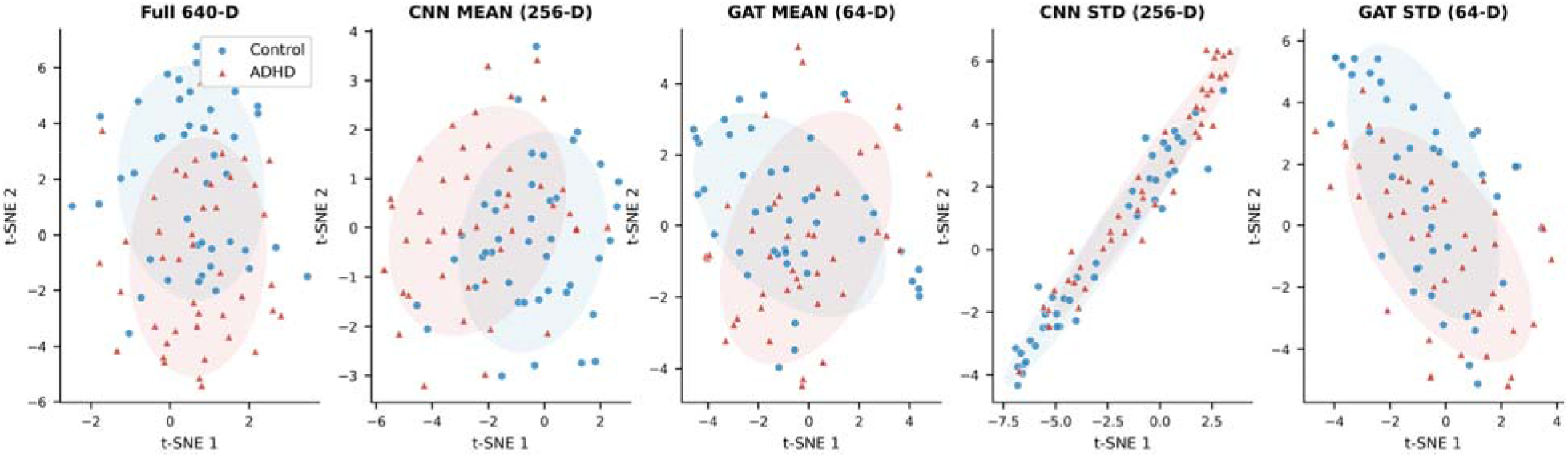
Two-dimensional visualization of ADHD and control representation geometry within the patient-level latent spaces. Each point represents one participant after averaging their out-of-fold patient-level embedding across the five cross-validation seeds (*N* = 86; 44 ADHD and 42 controls). Blue circles indicate control participants and red triangles indicate participants with ADHD. From left to right, the panels show the complete 640-dimensional ST-GAT embedding, CNN_mean_ (256 dimensions), GAT_mean_ (64 dimensions), CNN_std_ (256 dimensions), and GAT_std_ (64 dimensions). Shaded ellipses provide a descriptive summary of within-group dispersion in the two-dimensional projection and do not constitute inferential confidence regions. *Abbreviations*: CNN, convolutional neural network; GAT, graph attention network; ST-GAT, spatiotemporal graph attention network; std, standard deviation.

### 3.7. ST-GAT extracts unique graph-derived variance beyond CNN features

To test whether the GAT’s discriminative contribution is genuinely unique or merely inherited from the CNN backbone, a MANCOVA (Wilks’ Λ) was performed with two covariate framings (Table 3).

**Table 3.** Unique group-discriminative contribution of GAT embeddings beyond CNN-derived representations. . Multivariate analysis of covariance was used to determine how much of the ADHD–control separation in the GAT_mean_ and GAT_std_ blocks remained after adjustment for CNN-derived representations. Mean and std denote mean and standard-deviation pooling across each participant’s trials. Unadjusted *d*² quantifies the multivariate group effect in the GAT block before residualization, and residualized *d*² quantifies the effect remaining after adjustment for the respective CNN representation. GAT-unique variance was calculated as 100 × residualized *d*²/unadjusted *d*²; CNN-inherited variance was calculated as 100 minus the GAT-unique percentage. Wilks’ Λ and the associated *p* value test the residual multivariate ADHD–control effect after covariate adjustment. *Abbreviations*: CNN, convolutional neural network; GAT, graph attention network; MANCOVA, multivariate analysis of covariance; std, standard deviation.

| GAT Block | Covariate Framing | D<sup>2</sup> (Uncorr.) | D<sup>2</sup> (Resid.) | Inherited % | GAT-Unique % | Wilks $\Lambda$ | P |
| --- | --- | --- | --- | --- | --- | --- | --- |
| MEAN GAT | Independent EEGNet | 2.33 | 0.82 | 64.70% | 35.30% | 0.7662 | $2.9 \times 10^{-4}$ |
| MEAN GAT | Internal CNN branch | 2.33 | 0.93 | 60.10% | 39.90% | 0.7373 | $6.9 \times 10^{-4}$ |
| STD GAT | Independent EEGNet | 2.44 | 0.89 | 63.50% | 36.50% | 0.7511 | $2.9 \times 10^{-4}$ |
| STD GAT | Internal CNN branch | 2.44 | 0.84 | 65.70% | 34.30% | 0.7594 | $2.1 \times 10^{-4}$ |

Under the internal CNN branch framing (the ST-GAT’s own backbone as covariate), the GAT contributed 39.9% unique variance for the mean block and 34.3% for the std block, with Wilks’ Λ = 0.737 (p = 6.9 × 10) and 0.759 (p = 2.1 × 10), respectively. Under the stricter independent EEGNet framing, the GAT-unique proportion was 35.3% (mean) and 36.5% (std), with Wilks’ Λ = 0.766 (p = 2.9 × 10) and 0.751 (p = 2.9 × 10). All four tests were significant at p < .001, confirming that the GAT extracts discriminative structure not efficiently captured by either its own backbone or a separately trained CNN.

The consistency of the GAT-unique fraction (∼35–40%) across both framings supports the claim that the ST-GAT’s performance advantage over EEGNet is grounded in the representation geometry of the graph-derived dimensions, not in model complexity or regularisation alone. The CNN-inherited portion (60–66%) indicates substantial overlap with CNN-extracted features, as expected when both models process the same raw signal; the non-trivial unique residual demonstrates that the spatiotemporal PLV-constrained graph routing captures interaction structure that purely convolutional processing misses.

### 3.8. Behavioural characterisation and neural–behavioural linkage

The ADHD group showed significantly lower NoGo correct-rejection rates (63.1 ± 16.9% vs 76.0 ± 15.8%; t = −3.61, p = .0005) and higher false-alarm rates (37.8 ± 17.4% vs 24.5 ± 16.7%; t = 3.21, p = .002), consistent with the inhibitory-control deficit profile. Go hit rate was also lower in ADHD (94.7 ± 6.5% vs 98.0 ± 2.2%; t = −3.15, p = .003). Go reaction-time variability (SD and coefficient of variation) was directionally higher in ADHD but did not reach significance (Go RT SD: p = .254; Go RT CV: p = .167), indicating that the behavioural intra-individual variability effect is detectable but weak at this sample size.

A preliminary discriminative-axis projection was computed for each neural component block onto the ADHD-vs-Control direction. All four blocks showed significant group separation on the discriminative axis (all p < 10), with the full 640-d embedding achieving a large effect (d = 1.03). Among sub-blocks, GAT_mean_ (d = 0.93) and CNN_mean_ (d = 0.89) showed the strongest individual contributions, followed by CNN_std_ (d = 0.73) and GAT_std_ (d = 0.60).

While the present study establishes robust group-level discriminative axes, directly mapping these patient-level embedding projections to individual behavioral metrics (such as NoGo false-alarm rates or Go reaction-time variability) represents a distinct research question. Consequently, the neural embedding findings reported here are presented as group-level discriminative profiles rather than individual-difference correlates of behavioral performance, which remain to be characterized in future investigations.

### 3.9. Synthesis: ADHD classification emerges from representation distillation, not spatial localization

Together, these results show that the ST-GAT improves ADHD classification not because it learns a simple group-specific spatial attention map (Section: ADHD and control trials do not differ in spatial attention), but because it transforms diffuse trial-to-trial EEG variability into compact, graph-derived patient-level representations. The attention analysis establishes stable, task-relevant spatiotemporal trajectories that are consistent with the known multistage organization of inhibitory control. The embedding analyses identify the patient-level representational geometry that separates ADHD from controls: the GAT compresses widespread but redundant CNN-derived variability features into a small set of non-redundant graph-derived dimensions carrying ∼35–40% unique diagnostic variance beyond what a CNN captures alone. This pattern reframes the ADHD-related EEG signal as trial-to-trial variability distilled into stable patient-level representations, rather than as a single group-specific regional biomarker.

## 4. Discussion

The present study demonstrates that incorporating neurophysiologically informed functional connectivity into a graph-attention framework enables the learning of patient representations that capture the distributed organization of EEG activity during response inhibition. While the proposed ST-GAT consistently improved patient-level ADHD classification compared with the matched EEGNet baseline[23] (the model achieved a mean accuracy of 75.3%, compared with 70.0% for the matched EEGNet baseline), its principal contribution lies in how it represents neural activity rather than the magnitude of the performance gain alone. More importantly, reproducible window-resolved attention profiles coexisted with an absence of corrected group differences in spatial attention, whereas the graph-derived embedding blocks showed pronounced ADHD-control separation beyond that contained in the CNN representations. This dissociation supports a representation-distillation account: diagnostic information was expressed less as a localized attention signature than in the transformation of variable trial-level EEG activity into compact patient-level geometry. By combining ERP-informed temporal windows [14], PLV-defined graph structure [33], and trial-to-patient transfer learning, ST-GAT provides a proof of concept for neurophysiologically informed and auditable representation learning in moderate-sample EEG data.

This distinction is central to the neurophysiological interpretation of the model. Response inhibition comprises a sequence of partially dissociable cognitive and neurophysiological processes [13,41,42], and canonical EEG signatures such as the N2 and P3 reflect overlapping contributions of perceptual selection, conflict monitoring, attentional updating, response cancellation, and subsequent evaluation rather than process-pure indices of inhibition [14,43–45]. ST-GAT was explicitly organized around this temporal structure by segmenting each trial into ERP-informed windows. The resulting attention maps revealed reproducible routing preferences across these windows, indicating that the model repeatedly converged on temporally structured profiles compatible with the multistage organization of NoGo processing. This provides neurophysiological face validity and makes the internal behavior of the model more auditable. However, the attention coefficients should not be equated with causal feature importance, neural generators, or electrophysiological biomarkers. Attention weights specify how the model routes and aggregates information but do not establish which inputs are necessary for a particular prediction [3]. Accordingly, the absence of corrected ADHD-control differences indicates that classification did not depend on a detectable group-specific redistribution of attention at the tested lobe-by-window resolution. It does not demonstrate neurophysiological equivalence between the groups or exclude differences at finer spatial, temporal, spectral, or source-resolved levels. The attention profiles therefore primarily support model auditability and neurophysiological plausibility, whereas the patient-level embeddings identify where group-discriminative structure resides [46].

The interpretation of these maps must also reflect the precise architecture of the model. In the present implementation, each graph node receives the same 256-dimensional trial-level CNN embedding together with a learned node-identity embedding. The graph nodes therefore do not contain electrode-local trial features, and greater attention to a node or edge should not be interpreted as stronger weighting of the locally recorded signal at that electrode. Instead, the attention patterns reflect how a global trial representation is transformed and routed through a sensor topology constrained by PLV and temporal-window identity. The graph-derived representation should consequently be understood as the outcome of a relational transformation induced by the architecture rather than as a direct estimate of patient-specific functional connectivity. This technical distinction strengthens the central interpretation: the GAT does not merely amplify local EEG features but reorganizes a high-dimensional trial representation according to a neurophysiologically motivated relational structure.

The embedding analyses provide the strongest evidence for this transformation. CNN-derived variability features showed widespread univariate ADHD-control differences, whereas the lower-dimensional GAT blocks exhibited strong multivariate group separation despite containing substantially fewer individually significant dimensions. Moreover, after adjustment for either the internal CNN branch or the independently trained EEGNet representation, approximately 35%-40% of the unadjusted multivariate group-separation effect, expressed as, remained in the GAT blocks. This retained effect indicates that the graph-derived representations contain group-separating structure that is not linearly absorbed by the corresponding CNN representations. This should not be interpreted as the percentage of diagnostic variance uniquely explained by the graph. Nevertheless, the convergence of the two covariate framings supports the conclusion that the GAT branch performs more than a simple re-expression of CNN-derived features. It transforms them into a compact relational representation in which ADHD-control differences become strongly multivariate. This is consistent with the broader neurophysiological heterogeneity of ADHD, in which cognitive-control alterations are unlikely to be captured by a single stationary electrophysiological marker or regional activation difference [5,47,48]. Rather, ADHD-related alterations appear distributed across processing stages, frequency bands, spatial configurations, and levels of trial-to-trial variability [10–12,20]. The present findings extend this view computationally by showing that group information was not concentrated in a single spatial attention pattern but emerged in the geometry of a representation derived across trials and temporal windows.

A related methodological implication concerns shortcut learning. Highly flexible models can achieve apparently promising performance by exploiting regularities that correlate with diagnostic labels without representing the phenomenon of interest [1,49]. This risk is particularly pronounced in clinical EEG datasets in which the number of trials is large relative to the number of independent participants and nuisance variables such as age, sex, recording quality, vigilance, artifact structure, or preprocessing characteristics may covary with diagnosis [2,50]. Parameters of backward or decoding models are therefore not automatically interpretable as representations of the underlying neurophysiological processes [3]. The PLV-based graph addresses this problem by introducing an explicit neurophysiological inductive bias. PLV captures phase-consistent relationships between distributed EEG signals [32,33], which are relevant to the coordinated neural processing underlying response inhibition [14,34–37]. Constraining message passing through this topology narrows the model’s solution space toward phase-synchrony-informed relational transformations, while the transfer-learning procedure first exploits the relatively large number of trials to learn task-related representations and subsequently adapts these representations to the smaller number of patient-level observations. These design choices do not, however, demonstrate resistance to shortcut learning. They establish a neurophysiologically motivated and explicitly testable architecture. Controlled comparisons with alternative graph structures and direct tests of whether demographic or technical variables can be decoded from the learned embeddings will be required to determine whether the PLV prior specifically reduces shortcut reliance.

The findings also refine how intra-individual variability should be conceptualized in computational models of ADHD. ADHD has repeatedly been associated with increased behavioral and neural trial-to-trial variability [15–17], whereas conventional ERP averaging suppresses much of this variation to increase signal-to-noise ratio [19]. The observation that both mean- and standard-deviation-pooled representations carried diagnostic information should not be interpreted as evidence against the relevance of variability. Once the network has learned a task-relevant latent space, the mean location of a participant’s trials can itself reflect how variable trial-level activity has been transformed; a stable patient-level embedding can therefore be the product of variable neural processing rather than its opposite. At the same time, mean and standard-deviation pooling capture only the first two marginal moments of each participant’s trial distribution. They do not preserve covariance between latent dimensions, multimodality, temporal ordering, or transitions between trial states. The present results therefore show that trial-level variability contributes to patient representation, but they do not establish that ADHD is characterized by altered occupancy or transition dynamics of latent neural states. Future distributional, set-based, or state-space architectures could represent each patient as a distribution or trajectory of trial-level states and test whether ADHD involves altered state occupancy, persistence, or switching [51–53]. This would connect patient-level representation learning more directly to computational accounts of unstable cognitive-control states and metacontrol in ADHD [54].

The translational relevance of this framework lies less in producing another binary diagnostic label than in enabling a structured characterization of patient-level EEG organization. A clinically informative output could combine prediction confidence and calibration with data-quality indicators, window-resolved routing profiles, and the patient’s position relative to an independently validated reference distribution. Such outputs could support stratification, identify participants expressing a pronounced action-stopping-related EEG phenotype, and potentially quantify changes in latent representations across treatment or development. These applications will nevertheless require prospective validation and standardized EEG quality-control procedures before the framework can be interpreted as a clinical decision-support system [4,39]. The present dataset is moderate in size, and the diagnostic groups differed in sex distribution, age, and IQ. These variables represent potential confounds and should be examined through matching, residualization, or explicit confound-probe analyses. In addition, the ADHD sample is clinically heterogeneous. Although this increases ecological validity, it limits the extent to which the current model can distinguish effects associated with symptom dimensions, comorbidities, medication status, or diagnostic presentations.

Taken together, the findings establish ST-GAT as a proof of concept for neurophysiologically informed and auditable patient-level EEG representation learning. They suggest that ADHD-related information is expressed less as a localized spatial attention signature than in the way distributed trial-level activity is transformed into compact patient-level geometry. More broadly, the study illustrates a shift from evaluating clinical EEG-AI solely through classification accuracy toward examining the representations through which predictions are generated. Such models become scientifically more informative when their priors are explicit, their internal routing can be audited, and their learned patient spaces can be systematically tested against alternative representations and potential confounds.

## 5. Conclusion

This study demonstrates that neurophysiologically constrained graph attention can improve patient-level representation learning from EEG beyond conventional convolutional approaches. By integrating phase-synchrony-based connectivity with temporally resolved graph attention and patient-level aggregation, ST-GAT achieved improved classification between ADHD and control relative to EEGNet while providing insight into where discriminative information emerges within the learned representation. Importantly, the differences among two groups were not expressed as reliable group specific spatial attention patterns. Instead, the strongest separation emerged in the patient level latent space, where GAT-derived representations captured substantial discriminative information beyond both internal CNN features and the independent EEGNet baseline. Together, these findings demonstrate the potential of graph-based EEG models to capture clinically relevant neural dynamics within compact and interpretable patient-level representations.

## CRediT authorship contribution statement

**Rishabh Tiwari:** Conceptualization, Formal analysis, Methodology, Software, Visualization, Writing—original draft, Writing—review & editing, **Christian Beste:** Conceptualization, Funding acquisition, Project administration, Resources, Supervision, Validation, Writing—original draft, Writing—review & editing, **Negin Gholamipourbarogh:** Conceptualization, Project administration, Methodology, Supervision, Validation, Writing—original draft, Writing—review & editing.

## Declaration of generative AI use

During the preparation of this work, the author(s) used ChatGPT Pro (5.6 Sol) in order to improve clarity in writing. After using this tool/service, the author(s) reviewed and edited the content as needed and take full responsibility for the content of the published article.

## Acknowledgement

This work was supported by the Federal Ministry of Research, Technology and Space (Bundesministerium für Forschung, Technologie und Raumfahrt; BMFTR) as part of the German Center for Child and Adolescent Health (DZKJ) under the funding code 01GL2405B.

## Data availability

Materials availability: https://osf.io/47qun/overview

Data and code availability: https://osf.io/47qun/overview

